# Genetic circuits for feedback control of gamma-aminobutyric acid biosynthesis in probiotic *Escherichia coli* Nissle 1917

**DOI:** 10.1101/2023.06.09.544351

**Authors:** Matthew Lebovich, Lauren B. Andrews

**Affiliations:** University of Massachusetts Amherst, Department of Chemical Engineering, Amherst, MA, USA; University of Massachusetts Amherst, Biotechnology Training Program, Amherst, MA; University of Massachusetts Amherst, Molecular and Cellular Biology Graduate Program, Amherst, MA

## Abstract

Engineered microorganisms such as the probiotic strain *Escherichia coli* Nissle 1917 (EcN) offer a strategy to sense and modulate the concentration of metabolites or therapeutics in the gastrointestinal tract. Here, we present an approach to regulate production of the depression-associated metabolite gamma-aminobutyric acid (GABA) in EcN using genetic circuits that implement negative feedback. We engineered EcN to produce GABA by overexpressing glutamate decarboxylase (GadB) from *E. coli* and applied an intracellular GABA biosensor to identify growth conditions that improve GABA biosynthesis. We next employed characterized genetically-encoded NOT gates to construct genetic circuits with layered feedback to control the rate of GABA biosynthesis and the concentration of GABA produced. Looking ahead, this approach may be utilized to design feedback control of microbial metabolite biosynthesis to achieve designable smart microbes that act as living therapeutics.

## Introduction

Gamma-aminobutyric acid (GABA) is a non-essential amino acid produced by bacteria found in the human gastrointestinal tract (Strandwitz et al. 2019). It acts as a neurotransmitter that has been shown to affect neurological conditions including mood and sleep disorders (Diez-Gutiérrez et al. 2020; Sigel and Steinmann 2012) as well as epilepsy, anxiety, and depression (Diez-Gutiérrez et al. 2020; Kalueff and Nutt 2007; Ding et al. 2021). Therefore, it could prove beneficial to regulate GABA concentrations *in vivo*. Engineered probiotic bacteria such as *Escherichia coli* Nissle 1917 (EcN) have been proposed as a next-generation strategy to produce and modulate such gut metabolites. Currently, EcN has been used in clinical trials as a stand-alone therapeutic (Kruis 2004; Henker et al. 2007). Engineered EcN has been used to decrease metabolite concentrations, such as a strain developed to digest phenylalanine in patients with phenylketonuria (V. M. Isabella et al. 2018; V. Isabella et al. 2017), or to produce a variety of biomolecules (Amiri-Jami et al. 2015; Praveschotinunt et al. 2019; Forkus et al. 2017; Duan and March 2010). However, reliably controlling the concentration of a metabolite, such as GABA, *in vivo* requires synthetic regulation. Various strategies have been modeled and employed to introduce feedback and genetic circuits for cells to autonomously self-regulate the output level of a product in different cell types (Liu and Zhang 2018; Hu and Murray 2022; Saxena et al. 2016; Xie et al. 2016). Here, we sought to develop a model-guided approach to design feedback circuits to control biomolecular production in EcN using characterized genetic circuit components. For this work, we use the production of GABA as our test case and used a set of modular repressor-based NOT gates to construct feedback circuits (Nielsen et al. 2016; Stanton et al. 2014). These transcriptional NOT gates have been previously characterized in EcN by our group and allow us to build and model multiple feedback circuits easily.

To regulate the concentration of GABA *in vivo*, EcN cells need to both sense and produce it. Previously, we developed and characterized a GABA biosensor for EcN comprised of the *gabR* allosteric regulator from *Bacillus subtilis* 168 (Amidani et al. 2017) and a synthetic P_Gab_ promoter that is the transcriptional output signal of the sensor (Lebovich and Andrews 2022). Prior studies have engineered improved GABA production in *E. coli* by overexpressing the glutamate decarboxylases GadA and GadB, which convert glutamate into GABA, and overexpressing the glutamate/GABA antiporter GadC (Le Vo, Kim, and Hong 2012; Yu et al. 2019). In these studies, the amount of GABA produced was determined using high performance liquid chromatography (HPLC), which requires an extraction step (Kang et al. 2006) and cannot be done in a high-throughput manner. However, biosensors have been widely used for high-throughput screening to guide pathway optimization and metabolic engineering of products other than GABA (Rogers, Taylor, and Church 2016; Michener et al. 2012; Hossain et al. 2020).

Here, we use a biosensor-assisted approach to perform metabolic engineering of GABA biosynthesis in EcN and then integrate genetic circuits to regulate GABA production via feedback control. We perform genome editing to engineer EcN for GABA biosynthesis. Using a GABA biosensor, we screen the GABA production of these strains, and we also investigate the effects of pH and culturing conditions on GABA biosynthesis by these engineered EcN strains. Lastly, we construct and assay open loop and feedback genetic circuits for regulation of GABA production by EcN. Using modular, genetically-encoded NOT gates to construct feedback circuits, we were able to monitor and predictably regulate the rate of GABA production (Figure 1).

**Figure 1:**
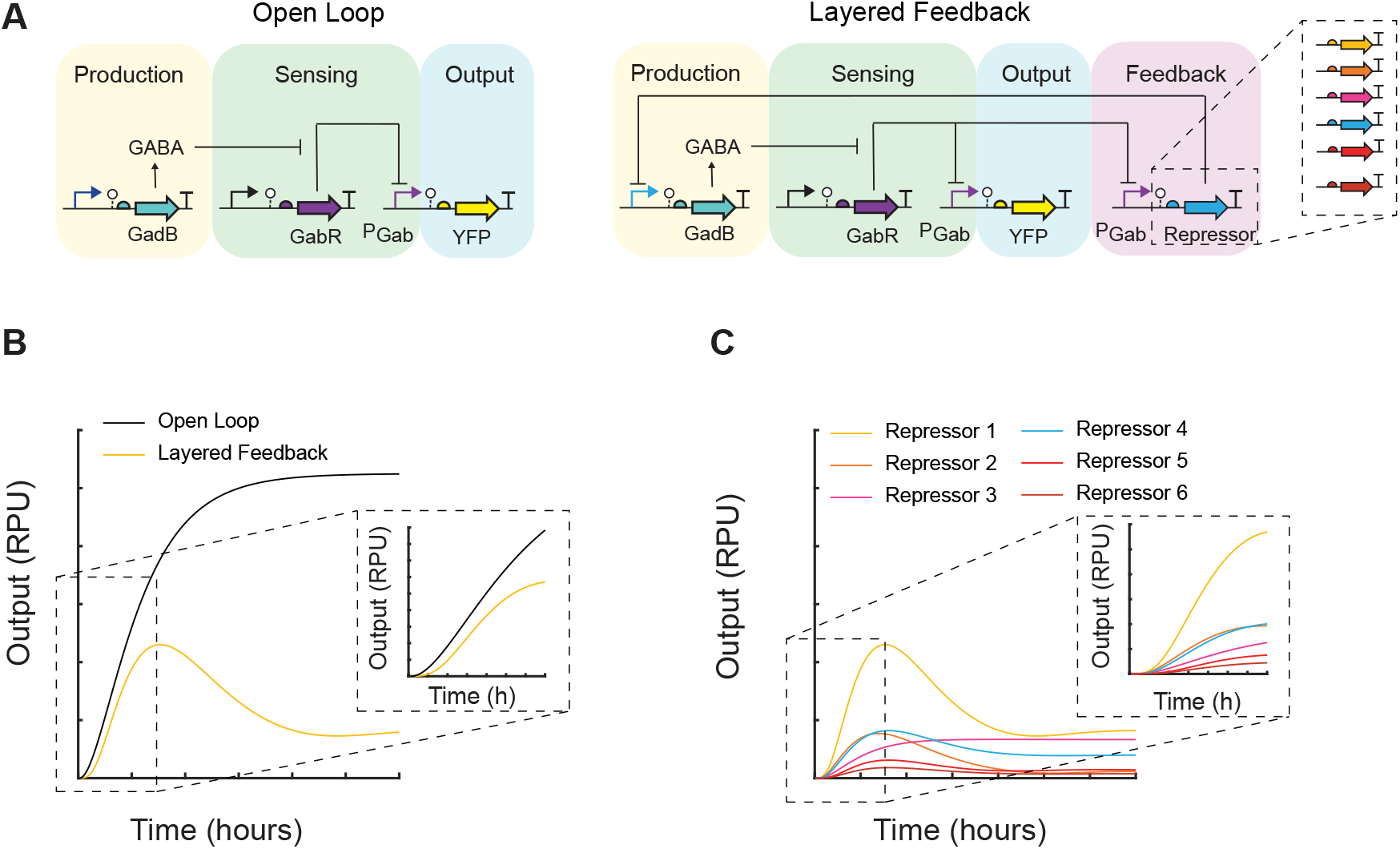
Theoretical framework for self-regulation of GABA biosynthesis in EcN with designable feedback. **(A)** The proposed open loop and layered feedback regulatory networks for GABA production are shown. In the open loop production module, expression of the enzyme GadB is induced and subsequently produces GABA. In the sensing module, GABA induces the P_Gab_ promoter of the GABA biosensor regulated by GabR. The P_Gab_ promoter in turn is the transcriptional driver for the output module, such as a yellow fluorescent protein (YFP) reporter. In the layered feedback design, the sensor output drives expression of a repressor-based NOT gate that controls the expression of the GadB enzyme, creating a negative feedback loop. These transcriptional NOT gates are interchangeable, and ones having different input-output responses can control the dynamics of GABA production. Mathematical models were constructed to simulate the output (P_Gab_) of these regulatory networks (Methods). **(B)** Simulation results comparing the output of the open loop and a layered feedback design. **(C)** Simulation results comparing layered feedback designs containing six different repressor NOT gates. The output is reported in standard relative promoter units (RPU) (Methods).

## Results

### Effects of growth conditions on GABA production

We first aimed to examine how changes in gene expression affected GABA production in EcN. To detect GABA, the previously developed GABA biosensor for EcN was utilized for screening (Lebovich and Andrews 2022). A yellow fluorescent protein (eYFP) reporter was placed under the control of the P_Gab_ sensor output promoter for all constructs. In addition to containing the GABA sensor, the plasmid pML3021 expressed *E. coli* glutamate decarboxylase GadB (Le Vo, Kim, and Hong 2012) under the control of the inducible promoter P_Tet_ (Figure 2A). In this system, GadB converts L-glutamate into GABA (Le Vo, Kim, and Hong 2012), which then induces the P_Gab_ promoter. The plasmid pML3021 was transformed into wildtype EcN as well as a strain with the chromosomal *gabT* and *gabP* genes deleted (EcN Δ*gabTP*). Deletion of *gabT* and *gabP* has been shown in previous work to increase GABA production (Yu et al. 2019). GabT digests GABA into succinate semialdehyde and GabP imports GABA (Yu et al. 2019), which may be beneficial to delete if the end goal is to export GABA *in vivo* (Figure 2B). Both strains were grown in 20 ml flask cultures containing M9 media supplemented with 35 g/L monosodium glutamate (MSG). To induce GadB production, 2 ng/mL anhydrous tetracycline (aTc) was added to the appropriate samples. Samples were grown at 37°C for 7 hours after which an aliquot was taken every hour. Single cell fluorescence was measured via flow cytometry and converted into RPU (Methods).

**Figure 2:**
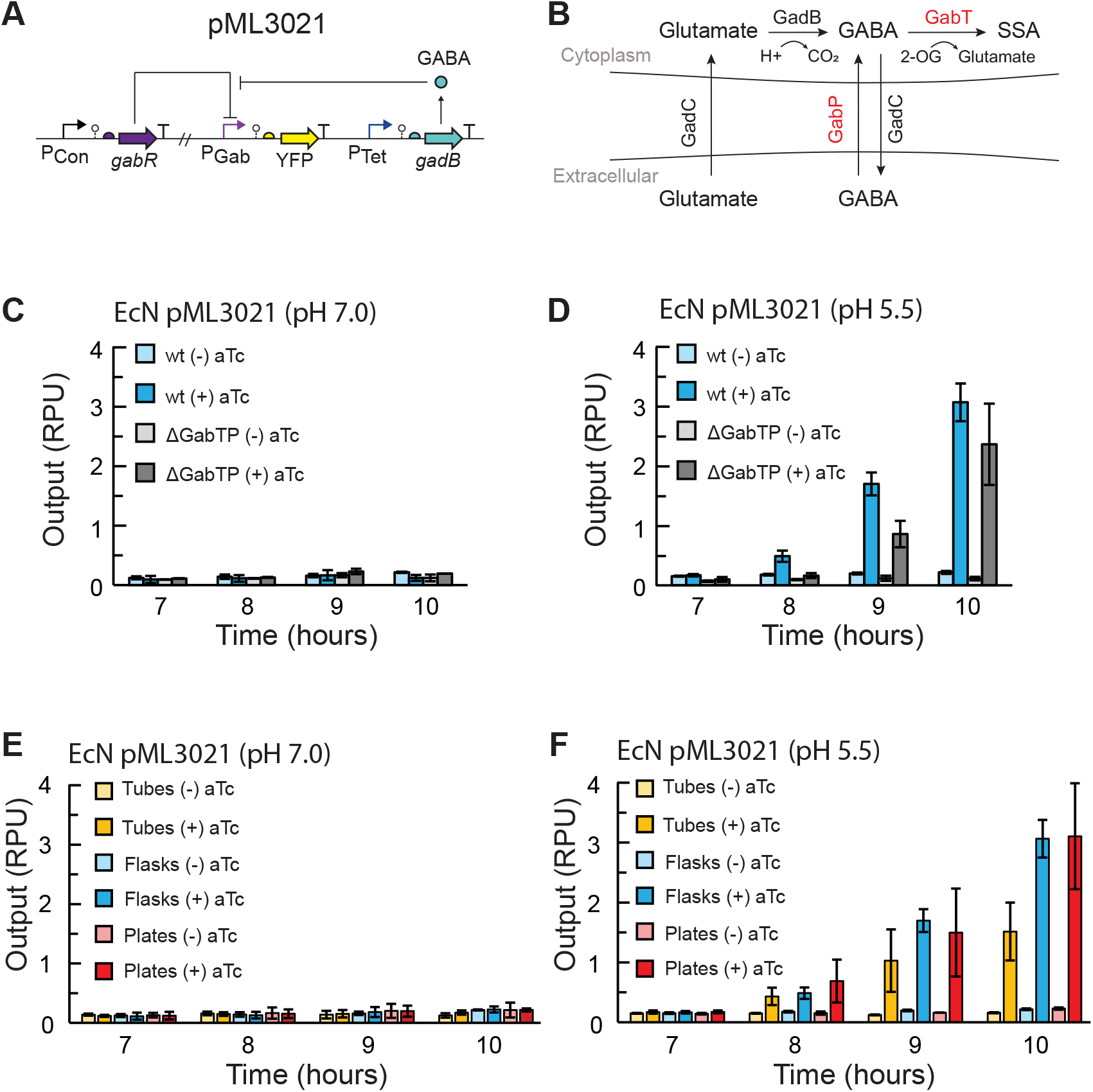
Using a GABA biosensor to screen strains and conditions for GABA production in EcN. **(A)** The GABA production plasmid pML3021 has inducible GadB expression. GadB produces GABA, which induces P_Gab_ and expression of eYFP via the GABA sensor regulated by GabR. **(B)** L-glutamate is transported into the cell via the antiporter GadC, and then converted into GABA by GadB. GABA can be degraded into succinate semialdehyde (SSA) via GabT. 2-oxogluterate (2-OG) is consumed in the reaction. GABA can also be imported into the cell by GabP. **(C)** GABA production assays for EcN without chromosomal editing (wt) and *ΔgabTP* chromosomal deletion strains harboring pML3021. Cells were grown in flask cultures without aTc (-) or with aTc added to a final concentration 2 ng/mL (+) at an initial pH = 7.0 or **(D)** initial pH 5.5. **(E)** GABA production assays for EcN wt harboring pML3021 grown in flasks, tubes, or plates without aTc (-) or with 2 ng/mL aTc (+) at an initial pH = 7.0 or **(F)** initial pH 5.5. All cultures were grown in M9 media supplemented with 35 g/L MSG and inoculated at an OD_600_ of 5×10^−5^. After 7 hours of growth, cell fluorescence was measured via flow cytometry at each time point. Fluorescence was converted into relative promoter units (RPU) (Methods). Bars represent the average of the measured median of a population of at least 10,000 cells assayed in three identical experiments performed on three separate days. Error bars represent the standard deviation.

When GadB was induced at pH 7.0, no significant change in P_Gab_ output was observed for any strains, which indicates negligible GABA production at all time points (Figure 2C). The output observed was equivalent to the basal activity of the P_Gab_ promoter, as previously reported for the sensor (Lebovich and Andrews 2022). This result is in agreement with previous literature reporting that GadB, which is notably part of an *E. coli* acid resistance system, is inactive at neutral pH (Pennacchietti et al. 2009; Capitani et al. 2003). When we lowered the initial medium pH to 5.5, we observed P_Gab_ activation beginning at 8 hours after induction with aTc, indicating GABA production (Figure 2D). In our experiments, deletion of *gabT* and *gabP* did not improve GABA production. To accurately determine the GABA concentration produced, we characterized the GABA sensor in each EcN strain in the identical growth conditions using exogenously added GABA (Supplementary Figure 1). When we accounted for the slightly different response of the sensor in each condition and strain, a comparable titer of GABA was produced with or without deletion of *gabT* and *gabP* and reached 0.053 ± 0.007 mM and 0.06 ± 0.01 mM, respectively, in M9 medium 10 hours after induction (Supplementary Figure 2). Given these results, we chose not to use the EcN Δ*gabTP* strain for subsequent experiments and used the wildtype EcN host instead.

We next examined how the culturing vessel used for growth affects GABA production. Previously, the GABA biosensor was assayed in small volume cultures using 96-well plates, which are commonly used for sensor characterization (Lebovich and Andrews 2022). However, this presented challenges in this work as there was an insufficient cell population to analyze multiple time points and cultures had low cell densities. To test how scaling up affected GABA biosynthesis, we performed the GABA production assay for the EcN pML3021 production strain in 96-well plates, 14-mL culture tubes, and 125-ml Erlenmeyer flasks. In media at pH 7.0, we again observed insignificant GABA sensor activation and negligible GABA production for all growth vessels (Figure 2E). At pH 5.5, we observed GABA sensor activation beginning 8 hours after GadB induction for cultures in plates, tubes, and flasks, and increasing activation through 10 hours (Figure 2F). Due to the very fast growth rate of EcN, we could not continue culturing beyond 10 hours and maintain exponential growth. When we used the GABA sensor response curves (Supplementary Figure 1) to determine the GABA concentration produced, interestingly, the amount of GABA is comparable and statistically indistinguishable for all three vessels (Supplementary Figure 3). However, the flask cultures had the highest reproducibility and lowest background GABA production without GadB induction. Therefore, we chose to use flask cultures in subsequent experiments.

### Feedback control of GABA production

After constructing our EcN GABA production strain, we next sought to further engineer this strain to have dynamic regulation of GABA biosynthesis via synthetic regulatory networks. To compare circuit topologies, we constructed genetic circuits with open-loop control (identical to our inducible GABA biosynthesis system) and layered-feedback control in which a negative feedback loop was created using a characterized repressor-based NOT gate, which acts as a transcriptional signal inverter (Figure 3A). In the latter design, the inducible promoter input expressing *gadB* (P_Tet_) was replaced by the repressible promoter P_Phlf_, which is the output promoter of the PhlF NOT gate (Stanton et al. 2014). The expression of the PhlF repressor was placed under the control of P_Gab_. In this way, the GABA sensor output feeds into the genetic circuit, which in turn regulates expression of GadB and GABA biosynthesis, creating the closed-loop feedback. GABA production of EcN cells transformed with the constructed plasmids were assayed in flasks using the conditions determined above and the GABA biosensor readout. We observed a slower rate of GABA production for cells containing the layered-feedback circuit as compared to the open loop (Figure 3B). This is consistent with predictions obtained from our mathematical models (Figure 3C) and in agreement with similar feedback loops reported in literature (Liu and Zhang 2018; Hu and Murray 2022). We observed detectable GABA production for both designs beginning 8 hours after inoculation. While we also predicted a lower concentration of GABA at steady state for the layered-feedback control, this could not be confirmed experimentally here using this batch growth system, given that growth could not be extended beyond 10 hours.

**Figure 3:**
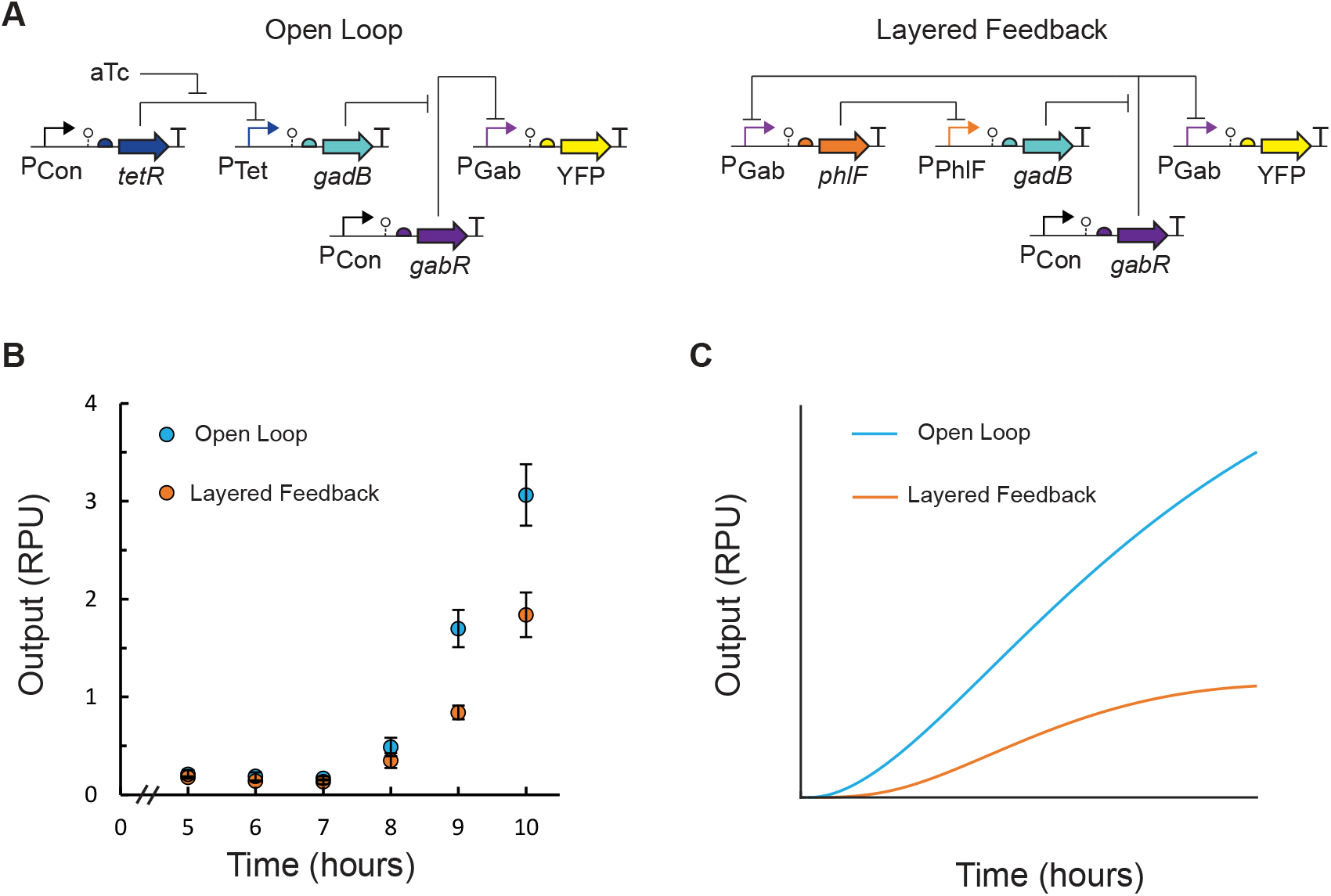
EcN GABA production controlled by open loop and layered feedback genetic circuits. **(A)** In the open loop control scheme (pML3021), GadB expression is induced by an exogenous inducer and produces GABA, which activates the P_Gab_ sensor output promoter of the GABA biosensor. The P_Gab_ signal is measured using an insulated eYFP transcriptional unit. In the layered feedback control scheme (pML3032), the P_Gab_ output of the GABA biosensor is the input to a transcriptional NOT gate (P_PhlF_ repressed by PhlF), which regulates GadB expression and GABA production detected by the GABA biosensor. **(B)** GABA production assays were performed for EcN cells containing plasmids pML3021 (blue) and pML3032 (orange). Cells were inoculated (OD_600_ = 5×10^−5^) in M9 media with MSG supplementation at pH 5.5 in 125 mL Erlenmeyer flasks. For cells containing the open-loop circuit, 2 ng/mL aTc inducer was added. After 5 hours of growth, cell fluorescence was measured via flow cytometry every hour. Fluorescence was converted into relative promoter units (RPU). Markers represent the average of the measured median of a population of at least 10,000 cells assayed in three identical experiments performed on three separate days. Error bars represent the standard deviation. **(C)** Simulation predictions of dynamic GABA production for the open loop (blue) and layered feedback (orange) designs from the corresponding mathematical model for each (Methods). Output is reported as the P_Gab_ sensor output in RPU.

Lastly, we posited that substitution of the repressor NOT gate in the layered-feedback circuit with another having different transfer function parameters could modulate the feedback and be used as a control strategy to achieve a desired rate of production and steady state output of GABA, as supported by modeling predictions (Figure 1C). For this experiment, we utilized a library of insulated repressor NOT gates (Nielsen et al. 2016) (Figure 4A). Across this set of six NOT gates, four repressors (PhlF, AmtR, IcaRA, and BM3R1) were chosen, and two gates contained an RBS variant for the repressor (PhlF, BM3R1) (Supplementary Table 3). We next constructed the 5 additional layered-feedback circuits, each containing a different NOT gate that has a unique response function in EcN (Figure 4B). EcN cells were transformed with each plasmid construct and assayed for GABA production (Figure 4C). As expected, we observed large differences in the rate of GABA production among the set of feedback circuits and up to 11-fold difference in output at 10 hours. We observed rough agreement with our predicted outputs from mathematical models containing the transfer function of each gate with the qualitative trends and rank order (Figures 1C and 4C).

**Figure 4:**
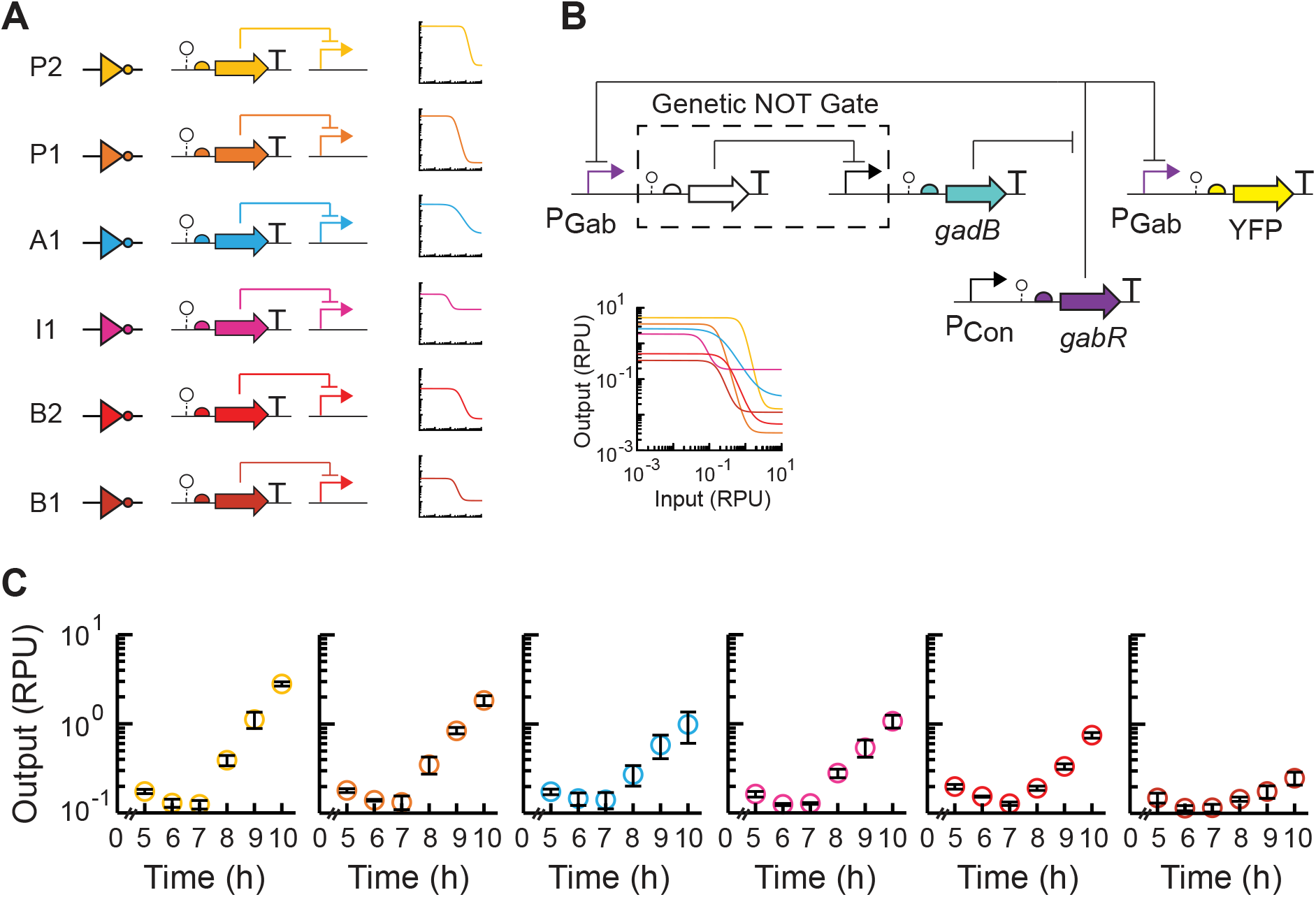
Tunable layered feedback using modular repressor NOT gates to control GABA production. **(A)** Six repressor-based NOT gates (Supplementary Table 3) each containing a TetR-family repressor and its cognate promoter were selected. The circuit NOT gate symbol, genetic schematic, and empirical response function in EcN is shown for each. Colors indicate the NOT gate identity. The overlay of the gate response functions is shown for comparison. **(B)** Each NOT gate was integrated into the closed-loop layered feedback circuit, and the corresponding plasmid designs were constructed. **(C)** GABA production assays were performed for EcN cells transformed with each feedback circuit plasmid. Cultures were inoculated (OD_600_ = 5×10^−5^) and grown in M9 supplemented with MSG at pH 5.5 in 125 mL Erlenmeyer flasks. After 5 hours of growth, cell fluorescence was measured via flow cytometry every hour. Fluorescence was converted into relative promoter units (RPU) of the P_Gab_ sensor promoter. Markers represent the average of the measured median of a population of at least 10,000 cells assayed in three identical experiments performed on three separate days. Error bars represent the standard deviation. Colors indicate the NOT gate identity shown in (A).

## Discussion

Engineered probiotic bacteria provide an opportunity to locally sense and control biomolecules in the gastrointestinal tract that would otherwise prove difficult to manipulate. Here, we utilize an intracellular sensor for the neurotransmitter GABA (Lebovich and Andrews 2022) to engineer and evaluate GABA production in EcN under various conditions, eliminating a rate limiting HPLC step (Le Vo, Kim, and Hong 2012). In this work, we found that deletion of *gabTP* did not improve GABA production and that physiologically relevant concentrations of GABA could be produced using only the GadB glutamate decarboxylase. While we found GadB expression alone to be sufficient for GABA production, this approach and the GABA biosensor could be applied for further metabolic engineering of GABA, such as by screening large libraries of GadB mutants or combinatorial pathways expressing GadA, GadB, and GadC (Yu et al. 2019). This work provides an additional demonstration of using biosensor-guided metabolic engineering, here shown in the clinically-relevant EcN bacterium.

Using feedback in gene regulatory networks, other researchers have been able to predictably and robustly control a range of cellular outputs (Liu and Zhang 2018; Hu and Murray 2022). Here, we build upon these works and implement closed-loop control via layered negative feedback to self-regulate GABA production in engineered EcN. We show that by utilizing an intracellular sensor and modular genetic circuit components, we can design and create feedback circuits to control GABA production and its dynamics. To understand the potential clinical viability of this approach, many further experiments are anticipated, such as understanding how expected *in vivo* perturbations affect feedback control. However, this work provides a step toward designing synthetic self-regulation of microbial metabolism for GABA biosynthesis in a gut microbe, and this approach may be broadly applied for *in vivo* control of metabolite biosynthesis in the gut.

## Materials and methods

### Strains, media, and inducers

*E. coli* Nissle 1917 was used for experimentally assaying all genetic constructs. *E. coli* NEB 5-alpha (New England Biolabs) and LB Miller medium (Fisher) was used for cloning. Cultures for assays, including overnight cultures, were grown in M9 media (Sigma-Aldrich; 6.78 g/L Na_2_HPO_4_, 3 g/L KH_2_PO_4_, 1 g/L NH_4_Cl, 0.5 g/L NaCl final concentration) with 0.34 g/L thiamine hydrochloride (Sigma-Aldrich), 0.2% w/v Casamino acids (Acros), 2 mM MgSO_4_ (Sigma-Aldrich), 0.1 mM CaCl_2_ (Sigma-Aldrich), and 0.4% w/v D-glucose (Sigma-Aldrich). Strains were assayed in M9 media with 35 g/L monosodium glutamate (MSG; Sigma-Aldrich). The pH was adjusted using hydrochloric acid (Fisher). The antibiotics used for selection were 50 μg/ml kanamycin (GoldBio), 100 μg/ml ampicillin (GoldBio), and 5 μg/ml chloramphenicol (GoldBio). The inducers used were anhydrotetracycline hydrochloride (aTc; Sigma-Aldrich) and gamma-aminobutyric acid (GABA; Sigma-Aldrich). GABA was stored as an aqueous solution, and aTc was dissolved in 100% ethanol.

### Construction of plasmids

The construction of the *gadB* containing plasmids was performed in two steps. First, part plasmids containing either a promoter or a multi-part construct (ribozyme, RBS, gene coding sequence, terminator) were combined into a transcription unit plasmid in a Type IIS DNA assembly reaction using BsaI-HFv2 (New England Biolabs). The destination vector (p15A origin of replication, ampicillin resistance) was supplied as a purified PCR product, and the genetic parts used were purified part plasmids. In the second Type IIS DNA assembly reaction using BbsI (New England Biolabs), these transcriptional unit constructs were assembled into the backbone plasmid pML3001 (Lebovich and Andrews 2022) containing kanamycin resistance, the p15A low-copy origin of replication and regulators for the sensors (*gabR* and *tetR*). The multi-part construct plasmids containing a ribozyme, synthetic RBS, *gadB*, and terminator were constructed by inserting them into a plasmid backbone containing kanamycin resistance and the colE1 ORI using a Type IIS DNA assembly reaction with the BbsI enzyme. The *gadB* gene was amplified from *E. coli* MG1655 genomic DNA. All parts and plasmids used in this work can be found in Supplementary Tables 1 and 2.

Type IIS DNA assembly reactions were performed in 5 μl total volume containing 20 fmol of each purified part or transcription unit plasmid, 10 fmol of the purified destination vector PCR product, 5 U of the appropriate Type IIS restriction enzyme, and 125 U T4 DNA ligase (2000 U/μl; New England Biolabs) in 1X T4 DNA Ligase Buffer (New England Biolabs). The reaction mixture was incubated in a thermal cycler (Bio-Rad C1000 thermal cycler, 105°C lid) with the protocol: 37°C for 6 hours, followed by 50°C for 30 min, and inactivated at 80°C for 15 min. Then, 2 μl of the assembly reaction was transformed into 5 μl chemically competent cells (*E. coli* NEB 5-alpha, New England Biolabs). Circuit constructs were analyzed by PCR. All transcriptional unit plasmids were sequenced by Sanger sequencing (Genewiz).

### Construction of gabTP deletion strain

The genes *gabT* and *gabP* were deleted from the EcN genome using the pSIJ8 plasmid (Jensen et al. 2016) containing the lambda Red recombineering system and ampicillin resistance (Addgene plasmid #68122). The plasmid was transformed into EcN via electroporation. Using the pKD3 plasmid (Datsenko and Wanner 2000) as the template, 500 bp homology arms (first 500 bp of *gabT* and last 500 bp of *gabP)* were added to either side of the chloramphenicol cassette to create pKD3-gab. Following the protocol previously described (Jensen et al. 2016), EcN cells harboring pSIJ8 were grown to an OD_600_ of approximately 0.3 in LB media (Fisher). The lambda Red proteins were induced by adding L-arabinose to a final concentration of 15 mM and growing the cells for an additional 45 minutes. The cells were then made electrocompetent and a purified PCR product of pKD3-gab containing the homology arms and chloramphenicol cassette was transformed into EcN harboring pSIJ8 via electroporation. The cells were recovered in SOC for 2 hours at 30°C and plated on an LB agar plate containing chloramphenicol and ampicillin. The plate was incubated at 30°C overnight. A single colony was streak purified at 30°C. To remove pSIJ8, the new strain was grown in liquid culture at 37°C, and a sample was streaked onto an agar plate with plate chloramphenicol. This last step was repeated until a colony from the chloramphenicol plate showed no growth on an ampicillin plate.

### GABA production assays

An overnight culture for each strain was started in M9 media (pH 7.0) from a freezer stock stored at -80°C. Cells were inoculated at an OD_600_ of 5×10^−5^ from an overnight culture and grown in M9 media with the appropriate antibiotics in 14 ml culture tubes (Fisher). Samples were grown in M9 media (pH 5.5 unless otherwise indicated) with 35 g/L MSG with appropriate antibiotics and inducers. Flask cultures were grown in 20 ml of M9 media in 125 ml Erlenmeyer flasks. Tube cultures were grown in 7 ml of M9 media in a 14 ml culture tube (Fisher). Plate cultures were grown in 200 μL of M9 media in the well of a 96-well U-bottom microtiter plate (Costar). Multiple wells from the same overnight were inoculated in plates to provide enough sample to analyze. All samples were grown at 37°C. Flasks and tubes were shaken at 250 rpm and 37°C in a shaking incubator (Innova 44R). Plates were incubated in an ELMI DTS-4 digital thermostatic microplate shaker at 1,000 rpm. After five hours of growth, a sample of each culture was taken every hour and incubated for 30 minutes at room temperature in PBS with 2 mg/ml kanamycin before fluorescence was measured using flow cytometry.

### Flow cytometry analysis

Cell fluorescence was measured using a BD Accuri C6 flow cytometer using a 480 nm blue laser and the FL1-A detection channel. The data for each sample was collected with a cutoff of 10,000 gated cell events at a flow rate of less than 1,000 events/s and at least 10,000 gated events collected per sample. For data analysis, the events were gated with a gate for cell-sized particles using FlowJo software. The median cell fluorescence for each sample was calculated using FlowJo. Histograms of cell fluorescence were generated using FlowJo. In all flow cytometry assays, the autofluorescence of wildtype EcN cells and the fluorescence of EcN cells containing the RPU standard plasmid were measured for samples from three separate colonies using identical dilutions and growth conditions as the assayed constructs.

The measured cell fluorescence in arbitrary units was converted to relative promoter units (RPU) as previously described (Nielsen et al. 2016) and using Equation 1 with the arbitrary unit values for the sample cell fluorescence (*YFP*), autofluorescence of wildtype EcN cells (*YFP*_*o*_), and fluorescence of EcN cells containing the RPU standard plasmid pAN1717 (Nielsen et al. 2016) (*YFP*_*RPU*_), which constitutively expresses eYFP. The sample fluorescence on each day’s experiment was converted to RPU. For each experiment, the average of the cell fluorescence from three separate colonies was used for the autofluorescence and RPU plasmid standard. The limit of detection was set to 0.001 RPU, and an output below this cutoff was set to this minimum value.

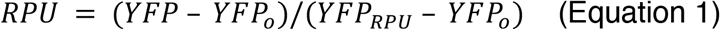

### Modeling

The following coupled differential equations were used to model the mRNA concentration of *gadB* (*M*_*B*_), concentration of GadB enzyme (*B*), concentration of GABA (*G*), transcriptional output of P_Gab_ (*μ*_*G*_), and transcriptional output of the promoter expressing *gadB* (*Q*_*r*_) and are based on a previously published model (Andrews, Nielsen, and Voigt 2018):

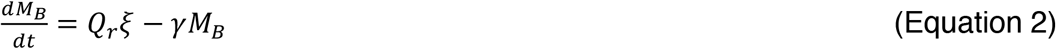

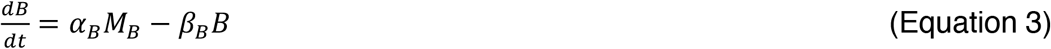

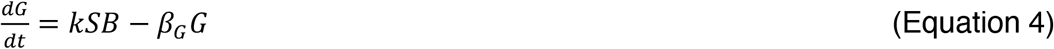

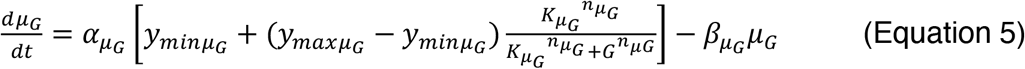

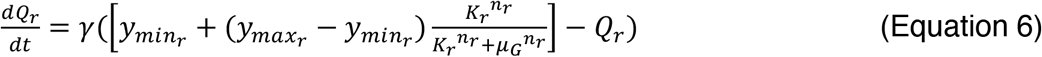

In the layered feedback circuits, *Q*_*r*_ is the output of the repressor NOT gate as shown in equation 6. In the case of the open loop regulatory network, *Q*_*r*_ was set to the maximum output of P_Tet_ and equation 6 was set to zero. The value of *μ*_*G*_ is the output of P_Gab_ in the GABA sensor. The parameters α and β represent the production and degradation rate constants of their respective species. The degradation rate of mRNA, *γ*, has been set to 0.025 min^-1^ as in previous work (Andrews, Nielsen, and Voigt 2018). A conversion factor *ξ* has been set to 0.025 [mRNA]min^-1^/RPU with *Q*_*r*_ = *M*_*Qr*_*ξ*^*-1*^*γ* (Andrews, Nielsen, and Voigt 2018). The parameters *y*_*min*_, *y*_*max*_, *K* and *n* are the values for the corresponding GABA sensor or NOT gate’s response function. Equations were solved using the ode45 function in MATLAB. Lists of all parameters used are in Supplementary Tables 3 and 4.

## Author Contributions

LBA and ML conceived of the study, designed experiments, and analysed data. ML performed the experiments. LBA and ML wrote the manuscript.

## Supplementary Material

The supplementary figures and supplementary tables are provided in the supporting information.

