## Supplementary Information for "Genetic circuits for feedback control of gamma-aminobutyric acid biosynthesis in probiotic *Escherichia coli* Nissle 1917"

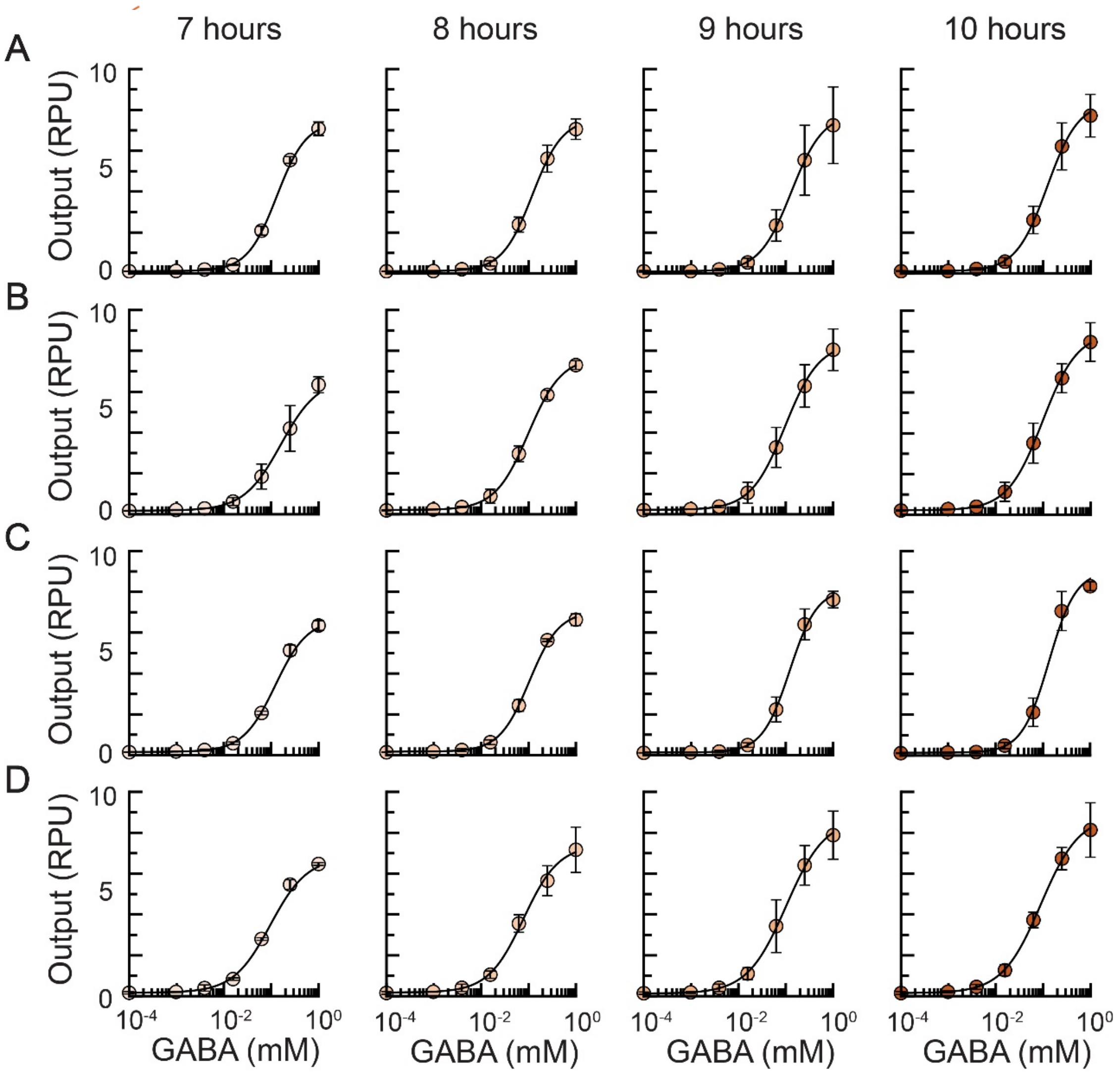

**Supplementary Figure 1: GABA sensor characterization in production conditions.** The GABA sensor characterization plasmid pML3009 was induced with exogenous GABA at each concentration and grown in the following strains and vessels. **(A)** EcN Δ*gabTP* pML3009 grown in flasks, **(B)** EcN pML3009 grown in flasks**, (C)** EcN pML3009 grown in culture tubes, and **(D)** EcN pML3009 grown in a 96-well microtiter plate. All cultures were inoculated at $OD_{600} = 5.0 \times 10^{-5}$. Single cell fluorescence was measured via flow cytometry (Methods). The markers represent the average of the median fluorescence measured on 3 separate days. The error bars are one standard deviation. The fitted response function for each is shown (solid line).

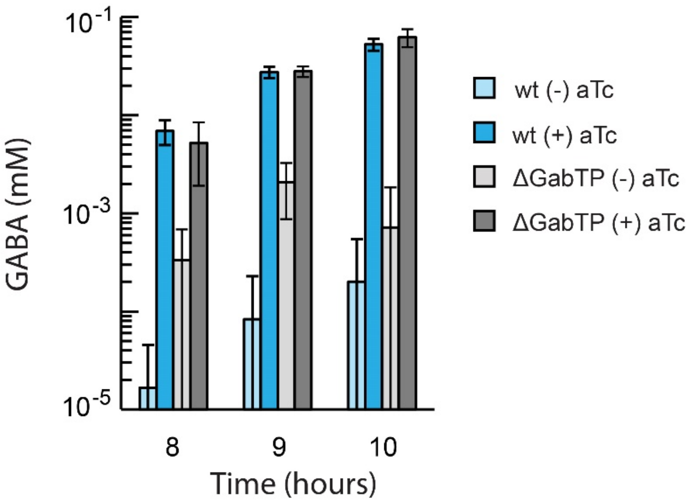

**Supplementary Figure 2: GABA production in EcN and EcN *ΔgabTP*.** For the data of the GABA production assays in Figure 2D, the output of the GABA biosensor was converted to GABA concentration using the corresponding fitted sensor response function (Supplementary Figure 1). The measured fluorescence from each trial was converted to GABA concentration. The bars represent the mean value from the 3 experiments and the error bars represent the standard deviation.

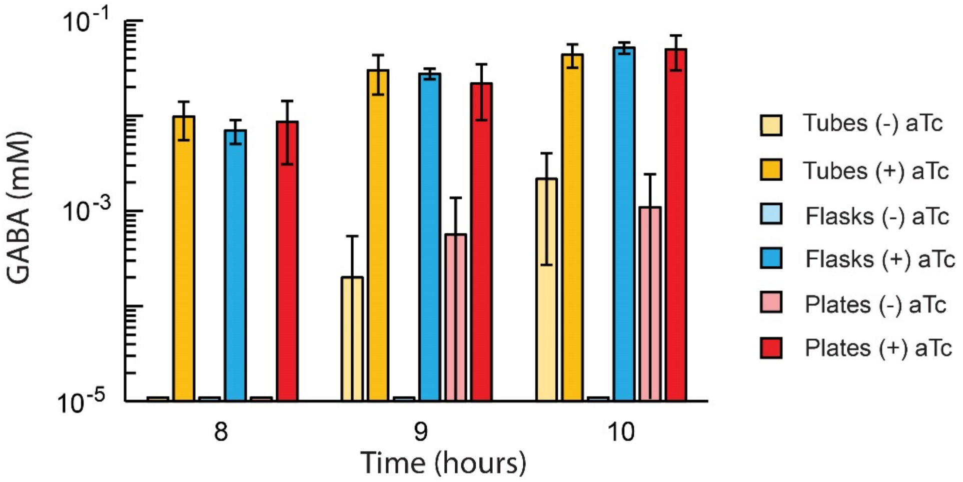

**Supplementary Figure 3: EcN GABA production in different growth vessels.** For the data of the GABA production assays in Figure 2F, the output of the GABA biosensor was converted to GABA concentration using the corresponding fitted sensor response function (Supplementary Figure 1). The measured fluorescence from each trial was converted to GABA concentration. The bars represent the mean value from the 3 experiments and the error bars represent the standard deviation. Values of 0 mM are displayed as 0.000011 mM on the plot.

### Supplementary Table 1: Genetic part sequences used in this work

| Part name | Type | DNA sequence | Source |
|---|---|---|---|
| $P_{AmtR}$ | Promoter | cttgtccaaccaaatgattcgttaccaattgacagtttctatcgatctat agataatgctagc | [1] |
| $P_{BM3RI}$ | Promoter | Aatccgcgtgataggtctgattcgttaccaattgacggaatgaacgt tcattccgataatgctagc | [1] |
| $P_{IcaRA}$ | Promoter | gtcaactcataagattctgattcgttaccaattgacaattcacctacct ttcgttaggttaggttgt | [1] |
| $P_{PhlF}$ | Promoter | cgacgtacggtggaatctgattcgttaccaattgacatgatacgaa acgtaccgtatcgttaaggt | [1] |
| $P_{Gab}$ | Promoter | ataccatcaaaaagttataattggtactttacggcataccaagtccta ggtactatgctagc | [2] |
| $P_{Tet}$ | Promoter | Tactccaccgttggctttttccctatcagtgatagagattgacatccc tatcagtgatagagataatgagcac | |
| A1 | RBS | aatgttccctaataatcagcaaagaggttactag | [3] |
| B1 | RBS | ctatggactatgttttaactactag | [3] |
| B2 | RBS | ctatggactatgtttttcaaagacgaaaaactactag | [3] |
| I1 | RBS | attgctatggactatgtttcaaagtgagaatactag | [3] |
| P1 | RBS | ctatggactatgtttgaaagggagaaatactag | [3] |
| P2 | RBS | ggagctatggactatgtttgaaaggctgaaatactag | [3] |
| *amtR* | Gene | atggcaggcgcagttggtcgtccgcgtcgtagtgcaccgcgtcgtg caggtaaaaatccgcgtgaagaaattctggatgcaagcgcagaa ctgtttacccgtcagggttttgcaaccaccagtacccatcagattgca gatgcagttggtattcgtcaggcaagcctgtattatcattttccgagca aaaccgaaatctttctgaccctgctgaaaagcaccgttgaaccga gcaccgttctggcagaagatctgagcaccctggatgcaggtccgg aaatgcgtctgtgggcaattgttgcaagcgaagttcgtctgctgctg agcaccaaatggaatgttggtcgtctgtatcagctgccgattgttggt agcgaagaatttgcagaatatcatagccagcgtgaagcactgacc aatgtttttcgtgatctggcaaccgaaattgttggtgatgatccgcgtg cagaactgccgtttcatattaccatgagcgttattgaaatgcgtcgca atgatggtaaaattccgagtccgctgagcgcagatagcctgccgg aaaccgcaattatgctggcagatgcaagcctggcagttctgggtgc | [1] |

| | | | |
|---|---|---|---|
| | | accgctgcctgcagatcgtgttgaaaaaaccctggaactgattaaacaggcagatgcaaaataa | |
| *bm3RI* | Gene | atggaaagcaccccgaccaaacagaaagcaatttttagcgcaagcctgctgctgtttgcagaacgtggttttgatgcaaccaccatgccgatgattgcagaaaatgcaaaagttggtgcaggcaccatttatcgctatttcaaaaacaaagaaagcctggtgaacgaactgtttcagcagcatgttaatgaatttctgcagtgtattgaaagcggtctggcaaatgaacgtgatggttatcgtgatggctttcatcacatttttgaaggtatggtgacctttaccaaaaaatcatccgcgtgcactgggttttatcaaaacccatagccagggcacctttctgaccgaagaaagccgtctggcatatcagaaactggttgaatttgtgtgcaccttttttcgtgaaggtcagaaacagggtgtgattcgtaatctgccggaaaatgcactgattgcaattctgtttggcagctttatggaagtgtatgaaatgatcgagaacgattatctgagcctgaccgatgaactgctgaccggtgttgaagaaagcctgtgggcagcactgagccgtcagagctaa | [1] |
| *icaRA* | Gene | gtgaaagacaaaattatcgataacgccatcaccctgtttagcgaaaaaggttatgacggcaccaccctggatgatattgcaaaaagcgtgaacatcaaaaaagccagcctgtattatcactttgatagcaaaaaaagcatctacgagcagagcgttaaatgctgtttcgattatctgaacaacatcatcatgatgaaccagaacaaaagcaactatagcatcgatgccctgtatcagtttctgtttgagttcatcttcgatatcgaggaacgctatattcgtatgtatgttcagctgagcaacacaccggaagaattttcaggtaacatttatggccagatccaggatctgaatcagagcctgagcaaagaaatcgccaaattctatgacgaaagcaaaatcaaaatgaccaaagaggacttccagaatctgattctgctgtttctggaaagctggtatctgaaagccagctttagccagaaatttggtgcagttgaagaaagcaaaagccagtttaaagatgaggtttatagcctgctgaacatctttctgaagaaataa | [1] |
| *phlF* | Gene | atggcacgtaccccgagccgtagcagcattggtagcctgcgtagtccgcatacccataaagcaattctgaccagcaccattgaaatcctgaaagaatgtggttatagcggtctgagcattgaaagcgttgcacgtcgtgccggtgcaagcaaaccgaccatttatcgttggtggaccaataaagcagcactgattgccgaagtgtatgaaaatgaaagcgaacaggtgcgtaaatttccggatctgggtagctttaaagccgatctggattttctgctgcgtaatctgtggaaagtttggcgtgaaaccatttgtggtgaagcatttcgttgtgttattgcagaagcacagctggaccctgcaaccctgacccagctgaaagatcagtttatggaacgtcgtcgtgagatgccgaaaaaactggttgaaaatgccattagcaatggtgaactgccgaaagataccaatcgtgaactgctgctggatatgattttggtttttgttggtatcgcctgctgaccgaacagctgaccgttgaacaggatattgaagaatttaccttcctgctgattaatggtgtttgtccgggtacacagcgttaa | [1] |
| *gabR* | Gene | atggatatcacgattacactcgatcgttcagaacaagccgattatatctatcagcaaatttatcaaaagctgaaaaaagaaatcctcagccg | [4] |

| | | | |
|---|---|---|---|
| | | caatctgctgccgcactcgaaggttccctccaagcgggagctggct<br>gaaaatctcaaggtcagcgtaaattcagtgaattcagcctatcagc<br>agctgctggctgaggggtatttgtacgccattgaacgaaagggtttc<br>ttcgtggaggaactagacatgttttccgccgaggagcaccctccatt<br>tgcactgccggatgacctaaaagagattcacatcgaccagagcg<br>attggatatcgttttcacacatgagttccgatacagaccattttccgat<br>caaaagctggttccgctgcgagcaaaaagcggcctcccgctcata<br>ccgcacgctcggcgatatgtcacatccgcaagggatatatgaagt<br>gagagcggccattacgaggctcatttccctgacgaggggtgtaaa<br>atgcaggccggaacaaatgatcataggggcaggcacacaggtg<br>ctcatgcagctgttgactgagcttttacccaaggaagccgtgtatgc<br>gatggaggagcctggctacaggcgcatgtatcagcttttgaagaat<br>gccggaaaacaagtaaagacgatcatgctggatgaaaaaggca<br>tgtcgattgctgaaatcaccagacagcagccagatgtgctggtgac<br>caccccgtcgcatcagtttccgtccggaacgattatgcctgtatcca<br>gaagaattcagctgctgaactgggcagccgaggagccgcgccg<br>atatatcattgaggacgattatgatagtgaattcacatatgatgtaga<br>cagtattccggcgctgcaaagcctcgaccgttttcaaaatgtcatct<br>atatgggaaccttttcaaagtcccttctccccggcttacggatcagct<br>atatggtgttgccgcctgagctgttgagggcatacaaacagcggg<br>gctatgatctgcagacttgctcatcactcacacagctcaccctgcag<br>gaatttatcgagtctggtgaatatcagaagcatataaaaaaaatga<br>agcagcattataaagaaaagagagaacgcctgatcaccgcttta<br>gaagcagagttcagcggagaggttaccgtaaaaggggcaaatg<br>cggggctgcattttgttaccgaatttgataccaggcgcaccgaaca<br>agacatcctgtcacatgctgccgggctgcagcttgaaatattcgga<br>atgagccgatttaacttgaaggaaaacaagcggcaaacgggca<br>ggcctgctctcattatcggctttgcacggctgaaggaagaagatatt<br>caggagggtgtgcagcggcttttcaaagcggtttacggacataaa<br>aaaatccccgttacaggggattga | |
| *eYFP* | Gene | atggtgagcaagggcgaggagctgttcaccggggtggtgcccatc<br>ctggtcgagctggacggcgacgtaaacggccacaagttcagcgt<br>gtccggcgagggcgagggcgatgccacctacggcaagctgacc<br>ctgaagttcatctgcaccacaggcaagctgcccgtgccctggccc<br>accctcgtgaccaccttcggctacggcctgcaatgcttcgcccgcta<br>ccccgaccacatgaagctgcacgacttcttcaagtccgccatgccc<br>gaaggctacgtccaggagcgcaccatcttcttcaaggacgacgg<br>caactacaagacccgcgccgaggtgaagttcgagggcgacacc<br>ctggtgaaccgcatcgagctgaagggcatcgacttcaaggagga<br>cggcaacatcctggggcacaagctggagtacaactacaacagc<br>cacaacgtctatatcatggccgacaagcagaagaacggcatcaa<br>ggtgaacttcaagatccgccacaacatcgaggacggcagcgtgc<br>agctcgccgaccactaccagcagaacacccccaatcggcgacgg<br>ccccgtgctgctgcccgacaaccactaccttagctaccagtccgcc | 5 |

| | | | |
|---|---|---|---|
| | | ctgagcaaagaccccaacgagaagcgcgatcacatggtcctgctggagttcgtgaccgccgccgggatcactctcggcatggacgagctgtacaagtaa | |
| *lacI* | Gene | atgaaaccagtaacgttatacgatgtcgcagagtatgccggtgtctcttatcagaccgtttcccgcgtggtgaaccaggccagccacgtttctgcgaaaacgcgggaaaaagtggaagcggcgatggcggagctgaattacattcccaaccgcgtggcacaacaactggcgggcaaacagtcgttgctgattggcgttgccacctccagtctggccctgcacgcgccgtcgcaaattgtcgcggcgattaaatctcgcgccgatcaactgggtgccagcgtggtggtgtcgatggtagaacgaagcggcgtcgaagcctgtaaagcggcggtgcacaatcttctcgcgcaacgcgtcagtgggctgatcattaactatccgctggatgaccaggatgccattgctgtggaagctgcctgcactaatgttccggcgttatttcttgatgtctctgaccagacacccatcaacagtattattttctcccatgaggacggtacgcgactgggcgtggagcatctggtcgcattgggtcaccagcaaatcgcgctgttagcgggcccattaagttctgtctcggcgcgtctgcgtctggctggctggcataaatatctcactcgcaatcaaattcagccgatagcggaacgggaaggcgactggagtgccatgtccggttttcaacaaaccatgcaaatgctgaatgagggcatcgttcccactgcgatgctggttgccaacgatcagatggcgctgggcgcaatgcgcgccattaccgagtccgggctgcgcgttggtgcggatatctcggtagtgggatacgacgataccgaagatagctcatgttatatcccgccgttaaccaccatcaaacaggattttcgcctgctggggcaaaccagcgtggaccgcttgctgcaactctctcagggccaggcggtgaagggcaatcagctgttgccagtctcactggtgaaaagaaaaaccaccctggcgcccaatacgcaaaccgcctctccccgcgcgttggccgattcattaatgcagctggcacgacaggtttcccgactggaaagcgggcagtga | [1] |
| *tetR* | Gene | atgtccagattagataaaagtaaagtgattaacagcgcattagagctgcttaatgaggtcggaatcgaaggtttaacaacccgtaaactcgcccagaagctaggtgtagagcagcctacattgtattggcatgtaaaaaataagcgggctttgctcgacgccttagccattgagatgttagataggcaccatactcacttttgccctttagaaggggaaagctggcaagattttttacgtaataacgctaaaagttttagatgtgctttactaagtcatcgcgatggagcaaaagtacatttaggtacacggcctacagaaaaacagtatgaaactctcgaaaatcaattagcctttttatgccaacaaggtttttcactagagaatgcattatatgcactcagcgctgtggggcattttactttaggttgcgtattggaagatcaagagcatcaagtcgctaaagaagaaagggaaacacctactactgatagtatgccgccattattacgacaagctatcgaattatttgatcaccaaggtgcagagccagccttcttattcggccttgaattgatcatatgcggattagaaaaacaacttaaatgtgaaagtgggtcctaa | [1] |
| *kanR* | Gene | atgagccatattcaacgggaaacgtcttgctccaggccgcgattaaattccaacatggatgctgatttatatgggtataaatgggctcgcgata | [6] |

| | | | |
|---|---|---|---|
| | | atgtcgggcaatcaggtgcgacaatctatcgattgtatgggaagcccgatgcgccagagttgtttctgaaacatggcaaaggtagcgttgccaatgatgttacagatgagatggtcagactaaactggctgacggaatttatgcctcttccgaccatcaagcattttatccgtactcctgatgatgcatggttactcaccactgcgatccccgggaaaacagcattccaggtattagaagaatatcctgattcaggtgaaaatattgttgatgcgctggcagtgttcctgcgccggttgcattcgattcctgtttgtaattgtccttttaacagcgatcgcgtatttcgtctcgctcaggcgcaatcacgaatgaataacggtttggttgatgcgagtgattttgatgacgagcgtaatggctggcctgttgaacaagtctggaaagaaatgcataagcttttgccattctcaccggattcagtcgtcactcatggtgatttctcacttgataaccttatttttgacgaggggaaattaataggttgtattgatgttggacgagtcggaatcgcagaccgataccaggatcttgccatcctatggaactgcctcggtgagttttctccttcattacagaaacggctttttcaaaaatatggtattgataatcctgatatgaataaattgcagtttcatttgatgctcgatgagtttttctaa | |
| BydvJ | Insulator | gggtgtctcaaggtgcgtaccttgactgatgagtccgaaaggacgaaacacccctctacaaataattttgtttaa | [3] |
| ElvJ | Insulator | gccccatagggtggtgtgtaccacccctgatgagtccaaaaggacgaaatggggcctctacaaataattttgtttaa | [3] |
| SarJ | Insulator | gactgtcgccggatgtgtatccgacctgacgatggcccaaaagggccgaaacagtcctctacaaataattttgtttaa | [3] |
| RiboJ53 | Insulator | gcggtcaacgcatgtgctttgcgttctgatgagacagtgatgtcgaaaccgcctctacaaataattttgtttaa | [3] |
| RiboJ64 | Insulator | aggagtcaattaatgtgcttttaattctgatgagacggtgacgtcgaaactccctctacaaataattttgtttaa | [3] |
| L3S2P55 | Terminator | ctcggtaccaaagacgaacaataagacgctgaaaagcgtcttttttcgttttggtcc | [7] |
| L3S2P24 | Terminator | ctcggtaccaaattccagaaaagacacccgaaagggtgtttttcgttttggtcc | [7] |
| ECK120015170 | Terminator | acaattttcgaaaaaacccgcttcggcgggtttttttatagctaaaa | [7] |
| L3S2P11 | Terminator | ctcggtaccaaattccagaaaagagacgctttcgagcgtcttttttcgttttggtcc | [7] |
| ECK120033737 | Terminator | ggaaacacagaaaaaagcccgcacctgacagtgcgggcttttttttcgaccaaagg | [7] |
| L3S2P11 | Terminator | ctcggtaccaaattccagaaaagagacgctttcgagcgtcttttttcgttttggtcc | [7] |

| | | | |
|---|---|---|---|
| ECK120033737 | Terminator | ggaaacacagaaaaaagcccgcacctgacagtgcgggcttttttttttcgaccaaagg | [7] |
| L3S3P21 | Terminator | ccaattattgaaggcctccctaacggggggcctttttttgtttctggtctccc | [7] |
| L3S3P51 | Terminator | aaaaaaaaaaaacaccctaacgggtgtttttttgtttctggtctccc | [7] |

**Supplementary Table 2: Plasmids used in this work**

| Plasmid name | Description | Source |
|---|---|---|
| pAN1717 | RPU strain expressing YFP under control of J23101 on a backbone expressing *lacI*, *tetR*, *kanR* and the p15a ori | [3] |
| pML3001 | Plasmid backbone containing g*abR*, *lacI* and *tetR* as well as the resistance gene *kanR* and the p15a ori. | [2] |
| pML3009 | Plasmid expressing YFP under the control of $P_{Gab}$ on the pML3001 backbone | [2] |
| pML3021 | Open loop GABA production circuit. $P_{Gab}$-YFP, $P_{Tet}$-GadB | This work |
| pML3030 | Feedback circuit using the IcaRA I1 NOT gate on the pML3001 backbone | This work |
| pML3031 | Feedback circuit using the AmtR A1 NOT gate on the pML3001 backbone | This work |
| pML3032 | Feedback circuit using the PhlF P1 NOT gate on the pML3001 backbone | This work |
| pML3033 | Feedback circuit using the PhlF P2 NOT gate on the pML3001 backbone | This work |
| pML3034 | Feedback circuit using the BM3RI B1 NOT gate on the pML3001 backbone | This work |
| pML3035 | Feedback circuit using the BM3RI B2 NOT gate on the pML3001 backbone | This work |

**Supplementary Table 3: Hill equation parameters for repressor NOT gates**

| Repressor | RBS | $y_{min}$ (RPU) | $y_{max}$ (RPU) | K (RPU) | n |
|---|---|---|---|---|---|
| AmtR | A1 | 0.032 | 2.597 | 0.209 | 1.881 |
| BM3R1 | B1 | 0.012 | 0.336 | 0.178 | 3.437 |
| BM3R1 | B2 | 0.005 | 0.517 | 0.317 | 2.865 |
| IcaRA | I1 | 0.187 | 1.847 | 0.066 | 3.772 |
| PhlF | P1 | 0.003 | 3.550 | 0.175 | 3.924 |
| PhlF | P2 | 0.015 | 5.306 | 0.843 | 4.880 |

**Supplementary Table 4: Model parameter values used**

| Repressor | Value | Units |
|---|---|---|
| $\alpha_B$ | 1 | [GadB]/([RNA]*min) |
| $\gamma$[6] | 0.025 | 1/min |
| $\beta_B$ | 0.1 | 1/min |
| $\beta_{\mu_G}$ | 20 | 1/min |
| $\xi$[6] | 0.025 | [mRNA]/(min*RPU) |
| $k$ | 0.25 | 1/min |
| $S$ | 35 | g/L |
| $\beta_G$ | 3 | g/(L*min) |
| $\alpha_{\mu_G}$ | 3 | RPU/min |
| $y_{min\mu_G}$[2] | 0.04 | RPU |
| $y_{max\mu_G}$[2] | 4.59 | RPU |
| $K_{\mu_G}$[2] | 16.23 | g/L |
| $n_{\mu G}$[2] | 0.90 | unitless |
